# Transposable elements modulate gene expression in response to oxidative stress in *Drosophila suzukii*

**DOI:** 10.64898/2026.09.08.750078

**Authors:** Miriam Merenciano, Zaïnab Belgaïdi, Corinne Régis, Lucille Gand, Ana Carolina Pires Das Dores, Cristina Vieira

## Abstract

Transposable elements (TEs) are major contributors to genomic and regulatory variation and can influence the expression of nearby genes in response to environmental stress. The invasive fruit fly *Drosophila suzukii*, whose genome is composed of nearly 50% TE-derived sequences, provides an excellent model to investigate the regulatory impact of TEs on stress responses. Although previous studies found no global association between TEs and oxidative stress transcriptomic response in this species, the effects of individual TE insertions remain largely unexplored.

Here, we investigated the role of two polymorphic intronic TE insertions, Mrp4-TE and inaE-TE, located within genes implicated in the oxidative stress response in *D. melanogaster*. Using wild-type strains of *D. suzukii*, we assessed the effect of these insertions on gene expression following oxidative stress induced by paraquat exposure. We found that their corresponding nearby genes *Mrp4* and *inaE* were upregulated after paraquat treatment and that this response was significantly stronger in flies carrying the corresponding TE insertions. Allele-specific expression analyses further showed that the alleles containing the TE insertions exhibited a tendency towards higher expression levels than alleles lacking the insertions, both under control and oxidative stress conditions, supporting a *cis*-regulatory effect. In addition, Mrp4-TE was associated with increased sensitivity to oxidative stress.

Together, our results provide the first evidence that TE insertions can modulate oxidative stress-responsive gene expression in *D. suzukii*. These findings highlight the importance of investigating individual TE insertions to understand the genetic and regulatory mechanism underlying stress responses, adaptation, and invasion success in this species.

## INTRODUCTION

Transposable elements (TEs) are repeated DNA sequences capable of moving within the genome (Bourque et al., 2018; Chuong et al., 2017; Wells & Feschotte, 2020). They are ubiquitous in eukaryotes and constitute a major source of structural and regulatory variation (Bourque et al., 2018; Casacuberta & González, 2013; Chuong et al., 2017; Wells & Feschotte, 2020). Although long considered mainly as genomic parasites, TEs are now recognised as important contributors to genome evolution and phenotypic diversity, occasionally providing raw material for adaptive responses to environmental challenges (Feschotte, 2023).

Stressful environmental conditions can influence TE activity and, conversely, TE insertions can affect the expression of stress-related genes (Capy et al., 2000; Horváth et al., 2017; Lanciano & Mirouze, 2018). Several studies have shown that specific TEs are transcriptionally activated under stress and may modulate nearby gene expression by the addition of transcription factor binding sites (TFBSs) or by modifying transcript structure, among others (de Oliveira et al., 2021; Horváth et al., 2017; Merenciano, Oliveira, et al., 2025). More specifically, in *Drosophila*, TE insertions have been associated with gene transcriptional responses to thermal, immune, xenobiotic, and oxidative stress (Guio et al., 2014; Mateo et al., 2014; Merenciano & González, 2023; Rech et al., 2019; Ullastres et al., 2015, 2021). Oxidative stress is a major physiological challenge linked to aging and environmental adaptation (Tatar, 1999). Insects, like other organisms, are subjected to oxidative stress upon exposure to herbicides and insecticides (Kodrík et al., 2015). Also, low temperature fluctuations and exposure to UV lead to oxidative stress (Kodrík et al., 2015). Thus, identifying TE insertions that influence oxidative stress responses provides valuable insight into how genetic and regulatory variation shapes stress tolerance.

The invasive fruit fly *Drosophila suzukii* offers a powerful model to investigate potential regulatory impact of TEs. Native to Southeast Asia, *D. suzukii* has rapidly spread across North America and Europe within less than a decade, demonstrating a remarkable ability to adapt to new environments (Fraimout et al., 2017). Recent genomic studies revealed that approximately 47% of the *D. suzukii* genome consists of TEs, with 75% of these insertions found at low frequencies, suggesting that they are recent and potentially active (Mérel et al., 2021). Notably, in this species, invasive populations from North America and Europe exhibited a higher TE load compared to native ones from Japan, which coincides with a reduced genetic diversity (Mérel et al., 2021). Although TE accumulation in the invasive populations could be explained by genetic drift, these findings would also support the hypothesis that TEs may help explain the paradox of invasion; the ability of species to successfully adapt to new environments despite the reduced genetic diversity likely caused by genetic bottlenecks during colonization events (Mérel et al., 2021). In changing environments, TE sequences can be co-opted as regulatory elements that modulate the expression of nearby genes. Moreover, stress-induced activation of TEs can generate new insertions, providing additional sources of genetic and regulatory variation that may contribute to adaptive potential (Horváth et al., 2017). However, no clear correlation has been found between TE content and bioclimatic variables in *D. suzukii*, suggesting a limited effect of environmentally induced TE activity (Mérel et al., 2021). Furthermore, a recent study showed that TEs are not transcriptionally activated following oxidative stress and that differentially expressed genes are significantly depleted of TEs in this species (Marin et al., 2021). However, although no global effect of TEs on gene regulation has been observed, specific TE copies may still exert regulatory effects following oxidative stress, as demonstrated in *D. melanogaster* (Guio et al., 2014; Mateo et al., 2014; Merenciano & González, 2023; Rech et al., 2019; Salces-Ortiz et al., 2020; Ullastres et al., 2015, 2021). Nevertheless, functional validation is needed to establish a direct link between these insertions, changes in the expression of nearby genes, and the resulting phenotypic outcomes.

In this work, we focus on two polymorphic TE insertions (Mrp4-TE and inaE-TE) located within the introns of the *Mrp4* and *inaE* genes, respectively, both of which are involved in oxidative stress responses in *D. melanogaster* (Huang et al., 2014; Huang & Haddad, 2007; Lin et al., 2014; Marin et al., 2021; Mérel et al., 2021; Monnier et al., 2002; Salces-Ortiz et al., 2020). We first characterized these insertions across wild-type populations of *D. suzukii* and assessed their impact on gene expression following oxidative stress induced by paraquat exposure. Using allele-specific expression (ASE) analysis, we further tested whether the presence of the TE affects expression of the associated allele. Our results showed that *Mrp4* and *inaE* increased their expression after paraquat treatment, and that this response was stronger in populations carrying the intronic TE insertions. Moreover, we found that Mrp4-TE and inaE-TE drove allele-specific expression changes after paraquat exposure, indicating a *cis*-regulatory effect of the insertion. Finally, we observed that the presence of Mrp4-TE was associated with increased sensitivity to oxidative stress.

Together, these findings demonstrated that particular TE insertions can modulate gene expression responses to oxidative stress in *D. suzukii*, adding to the growing evidence that TEs represent a relevant source of regulatory variation in wild-type populations. Furthermore, understanding the genomic mechanisms underlying stress responses provides valuable insights into the processes of adaptation and biological invasion.

## MATERIAL AND METHODS

### Fly stocks and rearing

*D. melanogaster* stocks were reared on nutritive fly food medium “LM” containing agar, maize flour, yeast, Nipagine, ethanol, and water, and kept in a controlled environment (25 °C, 60% relative humidity and 12:12 hour light/dark cycle). *D. suzukii* stocks were reared on “Dalton” medium containing agar, cornmeal, yeast, sugar, Nipagine, ethanol, and water, and kept in a controlled environment (22.5 °C, 70% relative humidity and 16:8 hour light/dark cycle).

#### D. suzukii wild-type populations

We used 33 *D. suzukii* strains collected in 2014 from six different locations spanning both native and invasive ranges: Sapporo (Japan, S), Tokyo (Japan, T), Montpellier (France, MT), Paris (France, L), Watsonville (USA, W), and Dayton (USA, SOK) (Table S1) (Marin et al., 2021; Olazcuaga et al., 2020). Isofemale strains were established from a single gravid female placed in a vial with a nutritive medium and maintained with a low larval density.

#### D. melanogaster CRISPR/Cas9 mutants

To generate *D. melanogaster* CRISPR/Cas9 mutants of the *Mrp4* and *inaE* genes, we used a coconversion/co-CRISPR strategy that aims at reducing the time and effort needed to identify CRISPR-induced changes due to a lack of an obvious phenotype and low frequency after editing (Ge et al., 2016; Kane et al., 2017). Briefly, we simultaneously targeted the gene of interest and a marker gene, *white*, which is an X-linked recessive gene that produces white-eyed flies when mutated. We used two previously validated guide RNAs (gRNAs) for the marker gene *white*: w-sgRNA-1 (5’-ataccattcctgctctttgg-3’), and w-sgRNA-2 (5’-gacaaccatttgaggtatac-3’) (Ge et al., 2016). To design gRNAs for the genes of interest, we first retrieved their genomic sequences from FlyBase (https://flybase.org/) (Öztürk-Çolak et al., 2024) and we aligned them to the *D. suzukii* ortholog sequence obtained from the SpottedWingFlyBase (http://spottedwingflybase.org/) (Chiu et al., 2013). We designed two guide RNAs (gRNAs) per target gene, ensuring a minimum separation of 50 bp between them: *Mrp4*-gRNA-1 (5’-atcctacctcggcaagcgga-3’), *Mrp4*-gRNA-2 (5’-gggcattacgagggctgcca-3’), *inaE*-gRNA-1 (5’-gtatgacatatcaagtgctc-3’), *inaE*-gRNA-2 (5’-ctgattagcgctgagacga-3’). gRNAs were designed using the flyCRISPR target finder (https://flycrispr.org/target-finder/) within coding regions that showed high homology between *D. melanogaster* and *D. suzukii*, and with no predicted off-targets. For each gene, the two target-specific gRNAs, along with the two gRNAs targeting the marker gene *white*, were cloned into the pCFD5 plasmid following the pCFD5 cloning protocol (https://crisprflydesign.org/wp-content/uploads/2023/12/pCFD5cloningprotocol.pdf) (Port et al., 2014). The pCFD5 plasmid containing the gRNAs was microinjected with a concentration of 350-400 ng/ul into approximately 300-350 embryos from the Cas9-expressing strain y[1] sc[*] v[1]; P{y[+t7.7] v[+t1.8]=nos-Cas9.R}attP2 (Bloomington stock number #78782) in the Drosophila Transgenesis facility of the Centro de Biología Molecular Severo Ochoa (CBM-CSIC, Spain). F0 emerged flies were crossed individually with a *white* mutant strain w[1118] (Bloomington stock number #3605). F1 females were screened for the white-eye phenotype. We did not obtain transformants for the *inaE* gene. For the *Mrp4* gene, white-eyed F1 females were backcrossed individually with the parental Cas9-expressing strain y[1] sc[*] v[1]; P{y[+t7.7] v[+t1.8]=nos-Cas9.R}attP2 for three generations to remove potential off-targets generated during the CRISPR/Cas9 editing process (Bassett & Liu, 2014; Merenciano et al., 2023; Merenciano & González, 2023; Port et al., 2020). In the second and third backcrosses, only red-eyed flies carrying the targeted mutation were selected for further crossing, to eliminate the *white* eye mutation from the genetic background. Then, red-eyed backcrossed flies were intercrossed until a strain homozygous for the target gene mutation was established. The presence of the targeted mutation in the *Mrp4* gene was checked in each generation by PCR using two primer combinations. The primer pair 5’-tggcaccaattgatagtacgtt-3’ and 5’-ttatggccaatgccaaggtt-3’ amplified a 2,258 bp fragment in the wild-type allele and a 490 bp fragment in the allele carrying the targeted deletion. The second primer pair 5’-gctgacggcacccaataaag-3’ and 5’-ttatggccaatgccaaggtt-3’ only amplified a 795 bp fragment in the wild-type allele. PCR bands suggesting the targeted deletion were confirmed by Sanger sequencing. Simultaneously, to ensure the same genetic background as the CRISPR/Cas9-generated mutant strains, the maternal Cas9-expressing strain y[1] sc[*] v[1]; P{y[+t7.7] v[+t1.8]=nos-Cas9.R}attP2 was initially crossed with the w[1118] strain. Red-eyed offspring were then backcrossed to the maternal Cas9-expressing strain for three successive generations to obtain the wild-type (WT) strain.

#### D. suzukii Mrp4-TE- and Mrp4-TE+ strains

To minimize the background effect on our experiments, we created homozygous strains for the presence (Mrp4-TE+) and homozygous strains for the absence (Mrp4-TE-) of the Mrp4-TE insertion starting from the Mrp4-TE heterozygous strain T20 (Tokyo, Japan) (Table S1). We first individually crossed virgin T20 females with T20 males. After allowing sufficient time for egg laying and confirming larval development on the fly medium, we individually screened by PCR all the parental flies for the presence or absence of the Mrp4-TE insertion with two primer pairs (see below). To establish the Mrp4-TE+ strain, we kept only vials in which at least one parent was homozygous for the Mrp4-TE insertion. Conversely, to generate the Mrp4-TE-strain, we kept only vials with at least one parent homozygous for the absence of the insertion. Then, we performed successive brother-sister crosses with emerged flies until we obtained flies homozygous for the presence of Mrp4-TE and homozygous for the absence. Final crosses were screened by PCR to confirm the presence or absence of the Mrp4-TE insertion.

### Identification of tandem repeats in *D. suzukii*

Tandem repeats within Mrp4-TE and inaE-TE insertions were discarded using *Tandem Repeats Finder* (TRF) with the following parameters: match = 2, mismatch = 3, indel = 5, match probability = 80, indel probability = 10, minimum alignment score = 20, and maximum period size = 15 (-h -d options) (Benson, 1999). The resulting TRF output file was parsed using a custom shell script to separate the results for each TE sequence into individual files. For each TE, the coordinates of detected tandem repeats were extracted, sorted, and merged using the *mergeBed* utility from *BEDTools* (Quinlan & Hall, 2010) to collapse overlapping or adjacent repeat regions. Finally, a custom Python script was used to compute the proportion of each TE sequence composed of tandem repeat regions, providing an estimate of the internal repeat content of each TE insertion. Following the approach of (Rech et al., 2022), insertions in which more than 80% of the sequence consisted of tandemly repeated regions were classified as potential tandem TEs. In our data, the repetitive content was below this threshold (20.1% for Mrp4-TE and 24.6% for inaE-TE), and therefore these insertions were not considered tandem repeats.

### Genotyping flies for presence or absence of Mrp4-TE and inaE-TE insertions in *D.suzukii*

To check the presence/absence of Mrp4-TE and/or inaE-TE, two pools of five-ten flies were screened by PCR with two different primer pairs per insertion, designed with the online software Primer-BLAST (Ye et al., 2012). Briefly, we designed a pair of primers flanking each TE insertion site (FL and R primers), which generate PCR products of different sizes depending on the presence or absence of the specific TE. Additionally, a primer within the TE sequence (I primer), used in combination with the R primer, produces a PCR product only when the TE is present (González et al., 2008). The primer pair Mrp4-TE-FL 5’-actttcacatacgtcacgcct-3’ and Mrp4-TE-R 5’-tttgcctcgaacgtaatgtgc-3’ amplified a 1,200 bp fragment in flies with the Mrp4-TE insertion and a 335 bp fragment in flies without it. The second primer pair Mrp4-TE-I 5’-gcaagcagcaaagcgagatt-3’ and Mrp4-TE-R 5’-tttgcctcgaacgtaatgtgc-3’ only amplified a 964 bp fragment in flies lacking Mrp4-TE. The primer pair inaE-TE-FL 5’-gggggcgaaacattgatgatg-3’ and inaE-TE-R 5’-aggaaaatggtagcgggaca-3’ amplified a 1,172 bp fragment in flies with the inaE-TE insertion and a 896 bp fragment in flies without it. The second primer pair inaE-TE-I 5’-acactacttagcatcgatttgcc-3’ and inaE-TE-R 5’-aggaaaatggtagcgggaca-3’ only amplified a 498 bp fragment in flies lacking inaE-TE.

### Gene expression analysis

#### Sample collection for D. melanogaster CRISPR/Cas9 mutants

90 five-seven day-old females from each CRISPR/Cas9 mutant strain were collected and placed in vials containing fresh “LM” medium, in groups of 20-25 individuals. For each sample, 20-25 guts from *Mrp4* mutant flies were dissected, flash-frozen in liquid nitrogen, and stored at -80 °C until further processing. This procedure was performed in three biological replicates. In parallel, the same protocol was applied to the WT strain.

#### Sample collection for D. suzukii wild-type populations and Mrp4-TE+ and Mrp4-TE-strains

The day before the experiment, 180 four-six day-old males and females from the S29, MT47, and W120 wild-type populations, and the Mrp4-TE+ and Mrp4-TE-strains were collected and placed in vials containing fresh “Dalton” medium, in groups of 30 individuals. The day of the paraquat exposure, flies were first starved for two hours by transferring them to empty vials. After this period, exposed flies were placed in vials containing fresh “Dalton” medium supplemented with 20 mM paraquat for 24 hours, while control flies were transferred to vials with regular “Dalton” medium. After that, for each sample, gonads were dissected out and discarded, and the remaining somatic tissue, fly carcasses, (20-30) were flash-frozen in liquid nitrogen, and stored at -80 °C until further processing. This procedure was performed in three biological replicates for each sex and/or biological condition (control or exposed to paraquat).

#### RNA extraction and cDNA synthesis

Samples were homogenized in QIAzol lysis reagent, and chloroform was added to separate the aqueous and organic phases. RNA was subsequently purified from the aqueous phase using the RNeasy Kit (Qiagen) according to the manufacturer’s instructions. Before RNA elution, an on-column DNase I treatment with the DNase TURBO^TM^ (Invitrogen) was performed for 30 minutes at 37 °C to eliminate genomic DNA contamination. cDNA was then synthesized from 150-1,000 ng of total RNA using the Reliance Select cDNA Synthesis Kit (Bio-Rad).

#### qRT-PCR analysis

*Mrp4* expression on *D. melanogaster* CRISPR/Cas9 and WT strains was measured using the forward primer 5’-gtccaacgatgtgggtcgat-3’ and the reverse primer 5’-atagcaacgccaaacatgga-3’. *Mrp4* expression on *D. suzukii strains* was measured using the forward primer 5’-tggatagcaccgttggaact-3’ and the reverse primer 5’-gcacgctggtccgctt-3’. *inaE* expression on *D. suzukii* strains was measured using the forward primer 5’-tggatgctgatggactaacgg-3’ and the reverse primer 5’-gcttttcaccgatccctcca-3’. All primer pairs showed high efficiencies (90-110%). Gene expression was normalized with the housekeeping gene *Rp49* (5’-cggatcgatatgctaagctgt-3’ and 5’-gcgcttgttcgatccgta-3’ primers). We performed the qRT-PCR analysis using the SsoAdvanced Universal SYBR Green Supermix (Bio-Rad) on a C1000 Touch Thermal cycler (Bio-Rad). Results were analysed using the difference between cycle threshold (dCT) method (Pfaffl, 2001).

### Allele-specific expression analysis (ASE)

For each TE analysed, we selected one strain homozygous for the presence of the TE insertions (S29) and one strain homozygous for their absence (W120), based on our PCR genotyping results. Both strains had been previously sequenced with long-read technologies (Marin et al., 2021). To identify single nucleotide polymorphisms (SNPs) linked to the presence of the TEs and located in the coding region of their nearby genes, we generated separate multi-FASTA files containing the coding sequences of each gene from both strains. These sequences were aligned using *ClustalW* (https://www.genome.jp/tools-bin/clustalw). Diagnostic SNPs identified in these alignments were confirmed by Sanger sequencing in the corresponding genomic regions in the S29 and W120 strains. We then performed approximately 30 crosses between ten S29 males and ten W120 virgin females to generate F1 heterozygous flies, in which ASE was measured. Groups of 30 males or females (five-seven day-old) from the F1 generation were exposed to 20 mM paraquat for 24 hours, while control flies were maintained under identical conditions without paraquat exposure, as described above. After that, flies were dissected and gonads were removed. RNA extraction and cDNA synthesis were performed as previously described, using three biological replicates per condition and sex. The cDNA samples were sent to an external company for primer design and pyrosequencing (EpigenDx, Inc.). Ratios between the allele containing the TE and the allele without the TE were calculated from the pyrosequencing data. To assess whether expression differs between alleles, ratios were log2 transformed, and a one-sample t-test was applied to test for deviation from zero.

### Transcription factor binding site detection

Genomic regions corresponding to *Mrp4* (NW_023496835.1: 3,496,240-3,503,148) and *inaE* (NW_023496845.1: 10,688,986-10,719,457) were defined using a BED file, based on the S29 genome assembly (Merenciano, Janillon, et al., 2025). For each gene, the region was extended to include 1 kb upstream of the transcription start site to capture the proximal gene regulatory region. To systematically scan these regions, sequences were partitioned into overlapping windows using *BEDTools* (Quinlan & Hall, 2010). Windows of 200 bp were generated with a step size of 150 bp (i.e., 50 bp overlap) using the *makewindows* function. The resulting windows were separated by genes based on scaffold identifiers and saved as independent BED files. DNA sequences corresponding to each window were extracted from the S29 genome assembly using the *getfasta* function, generating multi-FASTA files for *Mrp4* and *inaE*. To account for background nucleotide composition during motif scanning, a first-order Markov model was generated from the whole genome using the *fasta-get-markov* function from the MEME Suite (Bailey et al., 2015). The position weight matrix for the Cnc transcription factor (motif ID MA0530.1) was obtained from JASPAR in MEME format (Ovek Baydar et al., 2026). Motif scanning was performed using the FIMO tool from the MEME Suite (Bailey et al., 2015). Prior to scanning, multi-FASTA files were split into individual sequence files using a custom *awk* script. FIMO was then run independently on each sequence with a significance threshold of 1e-4 and the previously generated background model. For each gene, individual FIMO outputs were generated into a single table. Motif occurrences were considered significant with a q-value < 0.1 (Table S2).

### Survival assays

#### Preparation of fly vials

For paraquat survival assays, fly medium was supplemented with 10 mM paraquat (methyl viologen dichloride hydrate, Sigma-Aldrich, ref. 75365-73-0). Paraquat was added to the medium while it was cooling, prior to pouring it into the vials. For H_2_O_2_ survival assays, a round filter paper was placed at the bottom of the fly vial with 50 ul of 6% glucose solution containing 15% H_2_O_2_ (hydrogen peroxide solution 30 % (w/w) in H_2_O, with stabilizer, Sigma-Aldrich, ref. H1009-100ML).

#### Paraquat survival assays in D. melanogaster CRISPR/Cas9 mutants

The day before the experiment, 180 four-six day-old females from each CRISPR/Cas9 mutant and WT strain were collected and placed in vials containing fresh “LM” medium, in groups of 10 individuals. The day of the survival assay, flies were first starved for two hours by transferring them to empty vials. After that, we transferred 150 flies in groups of 10 into vials with “LM” medium supplemented with 10 mM paraquat. Survival was monitored by recording the number of dead flies in every vial at least once daily until all individuals had died. To minimize microbial growth, flies were transferred to fresh vials every five-six days. A total of 15 replicate vials per strain were used for paraquat exposure (Table S3).

#### Paraquat survival assays in D. suzukii wild-type populations and Mrp4-TE+ and Mrp4-TE-strains and H_2_O_2_ survival assays in Mrp4-TE+ and Mrp4-TE-strains

120-130 four-six day-old males and females from the S29, MT47, and W120 wild-type populations and/or the Mrp4-TE+ and Mrp4-TE-strains were collected and placed in vials containing fresh “Dalton” medium, in groups of 10 individuals. The following day, for paraquat survival assays, after two hours of starvation in empty vials, we transferred 90-100 flies in groups of 10 into vials with “Dalton” medium supplemented with 10 mM paraquat. Simultaneously, 30 flies were transferred to vials with regular “Dalton” medium in groups of 10 as controls (Table S4 and S5). For H_2_O_2_ survival assays, after two hours of starvation in empty vials, we transferred 90-100 flies in groups of 10 into previously prepared vials with a filter paper soaked with 50 ul of 6% glucose solution containing 15% H_2_O_2_. Simultaneously, 30 flies were transferred to vials with a filter paper soaked only with 50 ul of 6% glucose solution in groups of 10 as controls (Table S6).

Survival was monitored by recording the number of dead flies in every vial at least once daily until all individuals had died. To minimize microbial growth, flies were transferred to fresh vials (containing the oxidative agent) every five-six days. A total of nine-ten replicates vials per sex and strain were used for paraquat exposure, while three replicate vials per sex and strain were maintained as controls.

#### Survival data analysis

Survival data were analysed separately for males and females using Kaplan-Meier survival analysis combined with a log-rank test for statistical comparison. Briefly, survival probabilities over time were estimated using the *survfit* function in R, based on event status (dead or alive). Kaplan-Meier survival curves were generated with the *ggsurvplot* function from the *survminer* R package to visually represent survival trends. Finally, to assess differences in survival between strains, pairwise log-rank tests were performed using the *pairwise_survdiff* function, with p-values adjusted for multiple comparisons using the Benjamini-Hochberg method.

## RESULTS

### Polymorphic TE insertions are present in the *Mrp4* and *inaE* introns

Based on recent literature, we selected two TE insertions located within *D. suzukii* genes whose functions are associated with oxidative stress responses in *D. melanogaster*: Mrp4-TE and inaE-TE (Huang et al., 2014; Huang & Haddad, 2007; Lin et al., 2014; Marin et al., 2021; Mérel et al., 2021; Monnier et al., 2002; Salces-Ortiz et al., 2020).

On the one hand, Mrp4-TE is a non-full-length 864 bp polymorphic Class II Rolling-Circle (RC) insertion located in the first intron of the *Multidrug response protein 4* (*Mrp4* or *dMRP4)* gene (Figure 1A). In a previous study, this insertion was present in a native population from Japan while it was absent in an invasive population from France (Marin et al., 2021). Moreover, the gene harboring the insertion was differentially expressed after oral exposure to the oxidative agent paraquat in one of the populations, suggesting a genotype-by-environment interaction (Marin et al., 2021).

**Figure 1.**
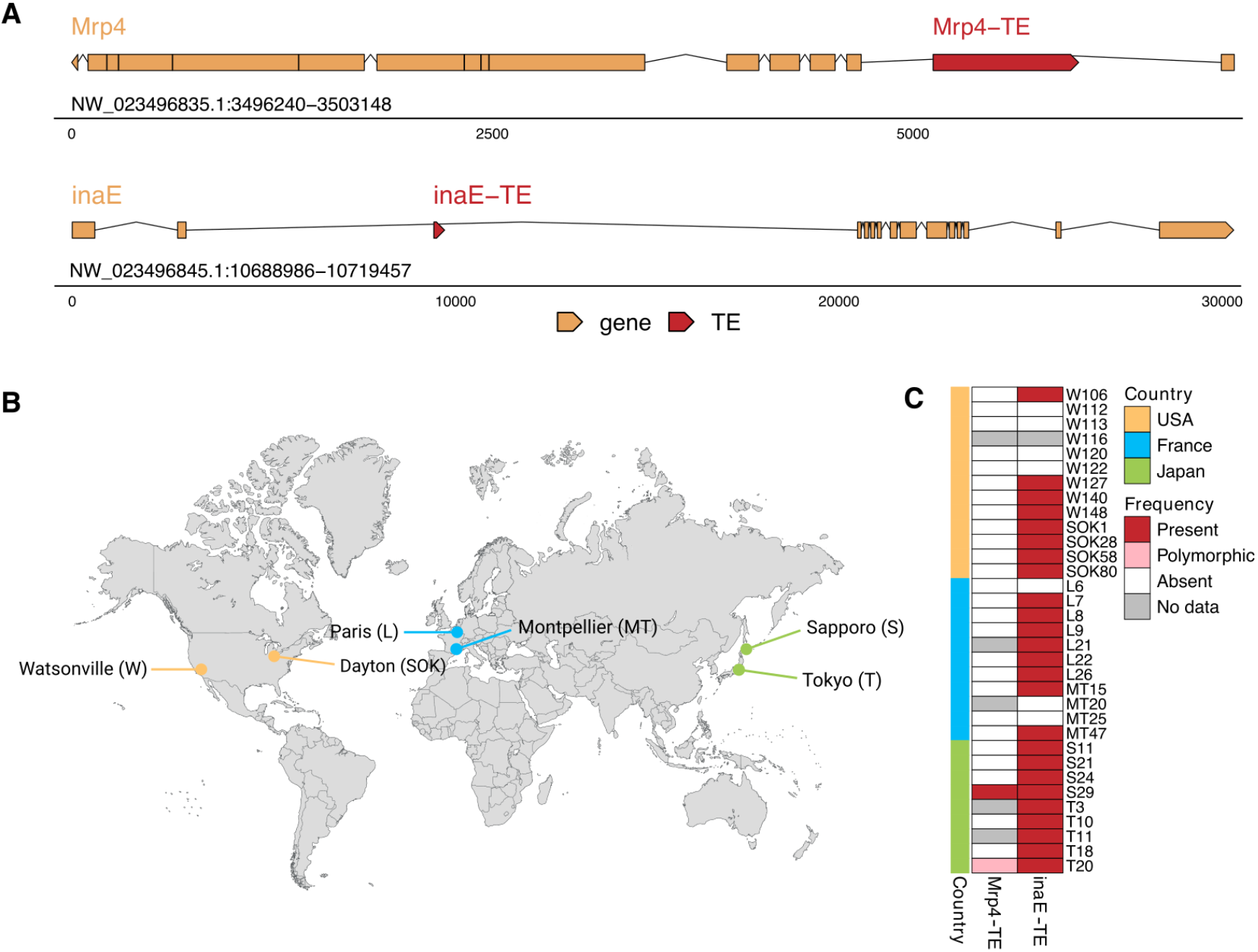
*Mrp4* and *inaE* genes harbour polymorphic TE insertions. **A**) Schematic representation of the genomic region where Mrp4-TE and inaE-TE are inserted in the S29 assembled genome. Orange boxes represent gene coding regions and untranslated regions (UTRs) while lines represent introns. TE insertions are represented as red boxes. **B**) Geographic origin of *D. suzukii* wild-type populations. Dots indicate the sampling locations of the populations analysed in this study. **C**) Heatmap showing the results of the PCR screening for the presence of the Mrp4-TE and inaE-TE insertions across strains. Red indicates strains homozygous for the TE insertion, pink indicates heterozygous strains, white indicates strains homozygous for the absence of the TE, and grey indicates undetermined genotypes.

On the other hand, inaE-TE is a non-full-length 276 bp polymorphic Class I LINE insertion located in the second intron of the *inactivation no afterpotential E* (*inaE)* gene (Figure 1A). This insertion was previously identified as a candidate adaptive element in a genome scan that combined three population-structure controlled methods implemented in the *BayPass* software (Mérel et al., 2021; Olazcuaga et al., 2020).

To discard the possibility that the Mrp4-TE and inaE-TE insertions correspond to tandem repeats, we analysed them with *Tandem Repeats Finder* (Benson, 1999). The proportion of repetitive regions within each insertion was below 80% (20.1% for Mrp4-TE and 24.6% for inaE-TE), and therefore they were not classified as tandem repeats (Rech et al., 2022).

We then assessed their presence in several wild-type populations of *D. suzukii*: two from the native range in Japan (Sapporo [S] and Tokyo [T]); two from invasive populations in France (Paris [L] and Montpellier [MT]); and two from invasive populations in the USA (Watsonville [W] and Dayton [SOK]) (Figure 1B). For each population, we assessed by PCR the presence or absence of the TE insertions in between four to nine isofemale strains. We found that Mrp4-TE was present in only one strain from the Sapporo population (S29) and was polymorphic in one strain from Tokyo (T20), but absent from all other strains (Figure 1C). Accordingly, its allele frequency was 0.2 in native Japanese populations and zero in the invasive populations from France and the USA. In contrast, inaE-TE was detected in 25 out of 32 strains tested and was present across all three countries, being fixed in Japan and with allele frequencies 0.73 in France, and 0.67 in the USA (Figure 1C).

### *Mrp4* and *inaE* genes are involved in oxidative stress response in *D. melanogaster*

The association of *Mrp4* and *inaE* genes with oxidative stress responses has already been studied in *D. melanogaster* (Huang et al., 2014; Huang & Haddad, 2007; Lin et al., 2014; Monnier et al., 2002; Salces-Ortiz et al., 2020).

*Mrp4* belongs to the MRP/ABCC subfamily of ATP-binding cassette (ABC) transporters. This gene has been associated with detoxification processes, as its protein product mediates the transport of a wide range of endogenous molecules and xenobiotics, including antiviral and anticancer drugs that can trigger oxidative stress and cellular toxicity (Borst et al., 2000; Dean & Annilo, 2005; Toyoda et al., 2008). In *D. melanogaster*, artificial overexpression of *Mrp4* increased sensitivity to oxidative stress induced by hydrogen peroxide (H₂O₂) and paraquat (Monnier et al., 2002). Furthermore, overexpression of this gene also delayed adult recovery from anoxia, suggesting a link between mechanisms that protect against both oxygen deprivation and oxidative stress (Huang & Haddad, 2007). Additionally, in wild-type flies exposed to paraquat, H₂O₂, or hyperoxic conditions, *Mrp4* expression levels were increased (Huang et al., 2014). In contrast, mutations causing reduced or negligible expression significantly reduced survival rates after acute paraquat exposure, while an overexpression was associated with the opposite phenotypic response (Huang et al., 2014). Thus, *Mrp4* expression has also been associated in promoting resistance to oxidative stress (Huang et al., 2014).

Since the expression of *Mrp4* has been linked to both sensitivity and resistance to oxidative stress, we sought to clarify its role by generating a homozygous *D. melanogaster Mrp4* knockout strain (Mrp4^-/-^) using the genome-editing CRISPR/Cas9 technique (see Material and Methods). qRT-PCR analysis confirmed a significant reduction in *Mrp4* expression levels (two-sided t-test, *p* = 0.025), and we observed that both mutant *Mrp4⁻/⁻* males and females showed reduced survival to an oral exposure to paraquat, confirming that *Mrp4* expression is involved in promoting resistance to oxidative stress in *D. melanogaster* (Log-Rank test, all *p* < 0.001) (Table S3 and Figure S1) (Huang et al., 2014).

*inaE* encodes a diacylglycerol lipase involved in phototransduction and response to oxidative stress. Indeed, mutant flies with reduced *inaE* expression showed a reduction in mean survival under paraquat-induced oxidative stress (Lin et al., 2014).

Altogether, these results led us to focus on *Mrp4* and *inaE* to investigate their potentially similar roles in *D. suzukii*.

### Paraquat-induced upregulation of *Mrp4* and *inaE* is stronger in wild-type *D. suzukii* populations carrying Mrp4-TE and inaE-TE insertions

To test whether the presence of the Mrp4-TE and inaE-TE insertions is associated with expression changes of their respective nearby genes, we quantified *Mrp4* and *inaE* expression in somatic tissues of males and females from three wild-type populations (S29, Japan; MT47, France; and W120, USA) under control and paraquat-induced oxidative stress conditions. Based on our previous results, Mrp4-TE was present in S29 flies, while inaE-TE was present in both S29 and MT47 flies (Figure 1C).

We found that paraquat exposure had a significant overall effect on *Mrp4* expression in both females and males (two-way ANOVA, all *p* < 0.001; Figure 2). Both the effect of strain (two-way ANOVA, females: *p* < 0.001; males: *p* = 0.002) and the interaction between strain and treatment were significant (two-way ANOVA, all *p* < 0.001; Figure 2), indicating that the magnitude of the paraquat response differs among strains (Marin et al., 2021). Post hoc analyses using *emmeans* revealed that paraquat induced a stronger response in S29 flies compared to MT47 and W120. In females, *Mrp4* expression increased approximately 12.3-fold in S29 (*p* < 0.001), 4.6-fold in MT47 (*p* < 0.001), and showed no significant change in W120 (0.3-fold, *p* = 0.766; Figure 2). A similar pattern was observed in males, with increases of approximately 12.8-fold in S29 (*p* < 0.001), 6.1-fold in MT47 (*p* < 0.001), and 3.3-fold in W120 (*p* = 0.007; Figure 2). These results suggested that the presence of the *Mrp4*-TE insertion could enhance *Mrp4* upregulation in response to paraquat.

**Figure 2.**
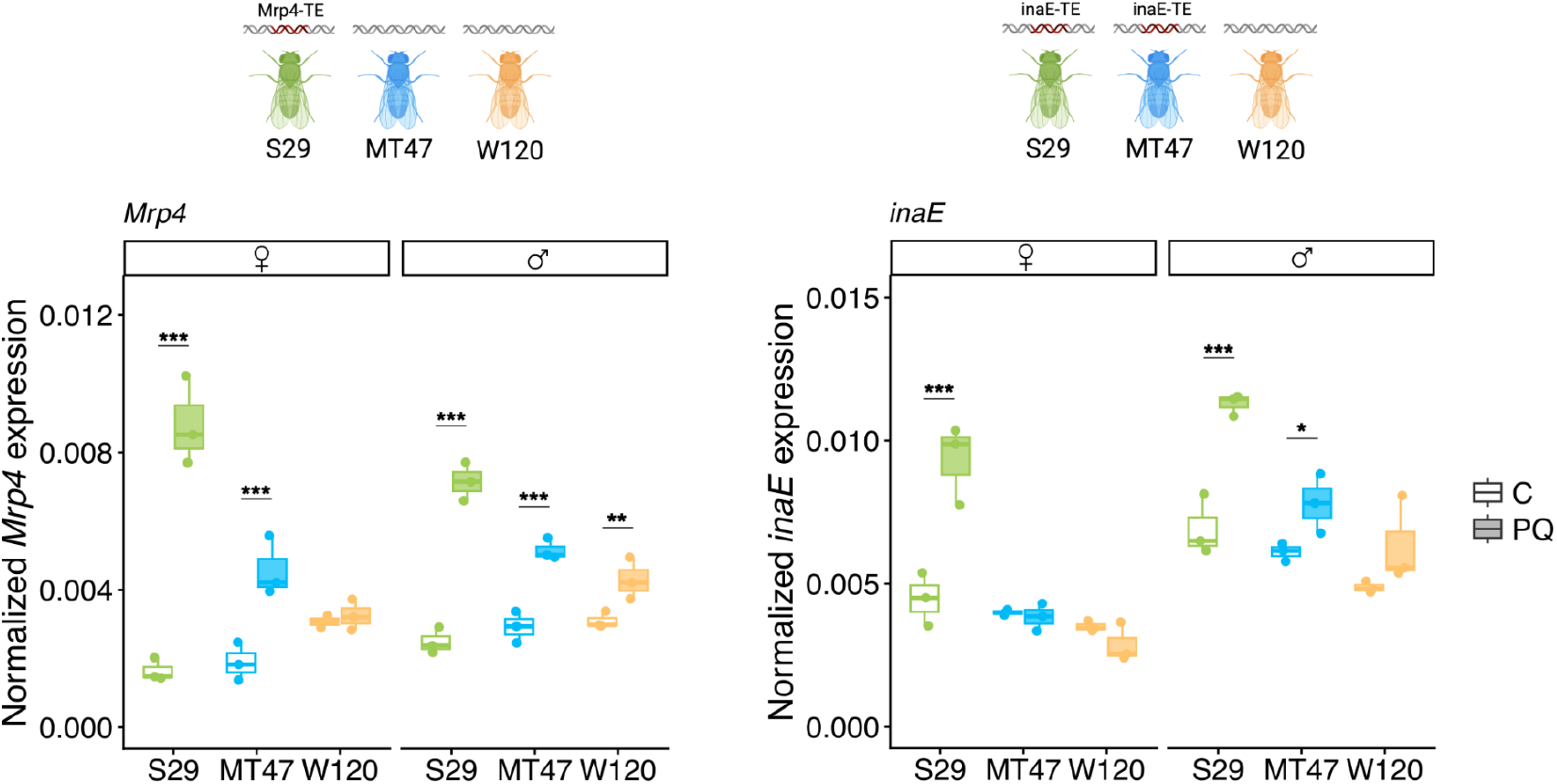
The *Mrp4*-TE and *inaE*-TE insertions are associated with stronger paraquat-induced upregulation of their respective genes in wild-type strains. Normalized expression of *Mrp4* (left) and *inaE* (right) with *Rp49* in S29, MT47 and W120 females and males, in control (C, empty boxes) and after paraquat exposure (PQ, filled boxes). Boxplot shows the median (horizontal line), first and third quartiles (lower and upper bounds, respectively), and minimum and maximum values (lower and upper whiskers, respectively). Asterisks indicate level of statistical significance (* p ≤ 0.05, ** p ≤ 0.01, ***p ≤ 0.001). Note that the fold changes shown in the figure are based on raw expression values, whereas the fold changes reported in the text are adjusted marginal means from the statistical model, which account for strain and treatment effects.

We also observed that paraquat exposure significantly affected *inaE* expression in both females and males (two-way ANOVA, females: *p* = 0.003; males: *p* < 0.001; Figure 2), with the response varying across strains. Both strain and the strain x treatment interaction factors were significant, confirming that *inaE* expression changes differed among fly strains (two-way ANOVA, all *p* < 0.05). As observed for *Mrp4*, S29 flies showed the strongest response. Indeed, in females, *inaE* expression increased approximately 7.8-fold (*p* < 0.001), whereas MT47 (-0.3-fold; *p* = 0.804) and W120 (-1.0-fold; *p* = 0.334) had no significant changes (Figure 2). In males, paraquat induced a 6.0-fold increase in S29 (*p* < 0.001), and smaller increases of 2.3- and 2.0-fold in MT47 and W120, respectively, which were statistically significant only for MT47 (*p* = 0.038; W120 *p* = 0.066; Figure 2). These results suggested that the presence of inaE-TE may be associated with increased *inaE* expression, at least in males, as paraquat-induced upregulation was observed in S29 and MT47, both of which carry the insertion, but not in W120, which lacks it. Therefore, it would also suggest that the putative regulatory effect of inaE would be background specific.

Overall, the presence of the *Mrp4*-TE and *inaE*-TE insertions appears to promote stronger paraquat-induced upregulation of their respective genes in wild-type strains, with *inaE* effects observed primarily in males and depending on the strain.

### Mrp4-TE and inaE-TE drive allele-specific expression changes after paraquat exposure

The previously found differences in *Mrp4* and *inaE* expression following paraquat exposure between wild-type strains could be due to multiple genomic differences. To further investigate whether Mrp4-TE and inaE-TE insertions influence the expression of their neighboring genes, we quantified allele-specific expression (ASE) in flies heterozygous for each insertion (see Material and Methods). Because both alleles coexist in the same cellular environment, any observed expression difference between them reflects functional *cis*-regulatory variation, likely driven by the presence or absence of the TE (Almlöf et al., 2014; Ullastres et al., 2021; Wittkopp et al., 2004).

To generate heterozygous flies, we crossed S29 individuals carrying both Mrp4-TE and inaE-TE insertions with W120 flies, lacking these insertions (Figure 3A, Figure 1C). We then measured *Mrp4* and *inaE* expression in the heterozygous offspring under both control and paraquat-treated conditions.

**Figure 3.**
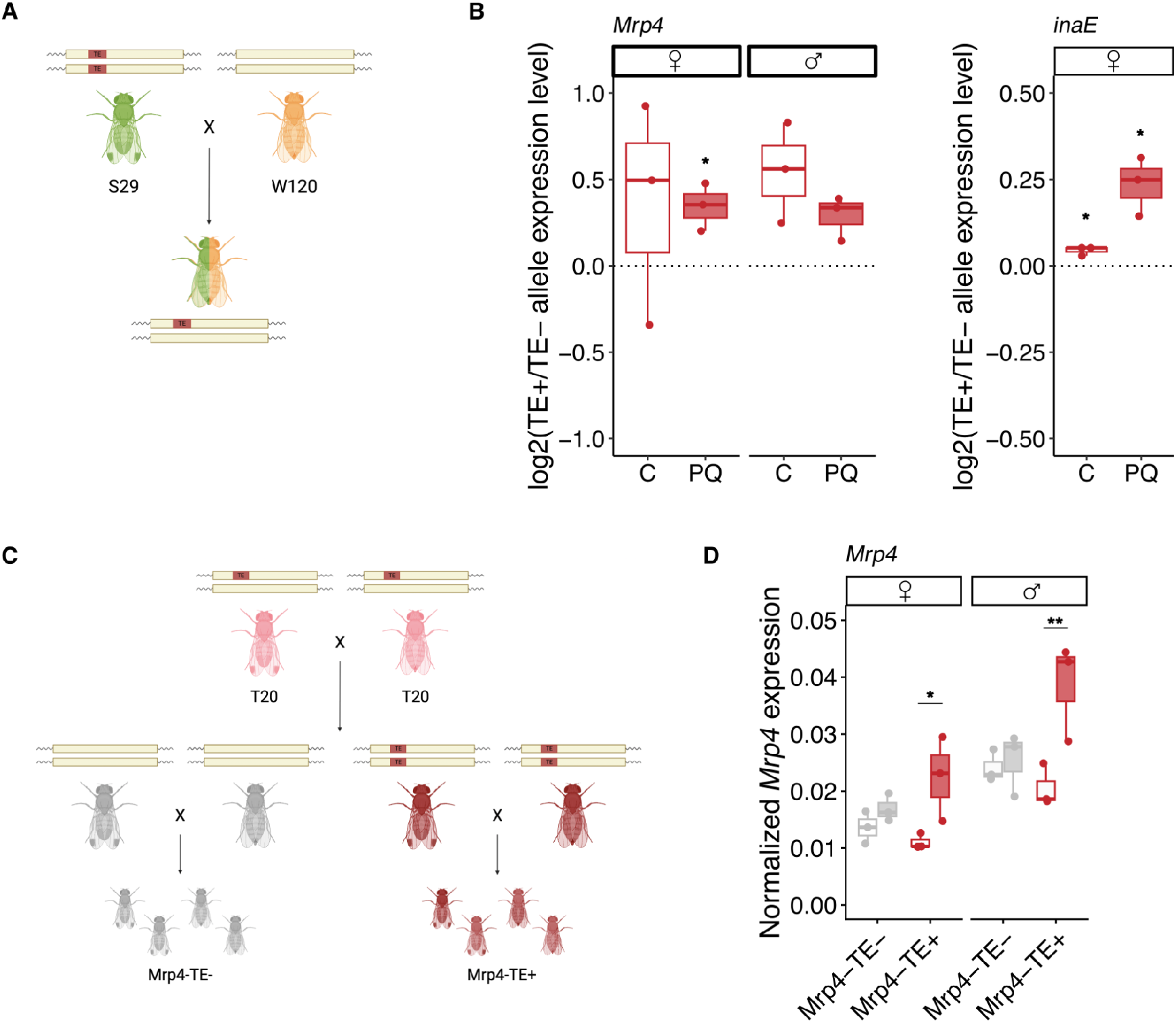
Mrp4-TE and inaE-TE insertions enhance paraquat-induced upregulation of their respective nearby genes. **A**) Schematic representation of the crosses used to generate heterozygous flies carrying both Mrp4-TE and inaE-TE insertions, in which the ASE experiment was performed. TE insertions are shown as red boxes. **B**) ASE results for both *Mrp4* (left) and *inaE* (right) genes in control (C, empty boxes) and after paraquat exposure (PQ, filled boxes). Boxplots represent the average ratio of gene expression levels between the allele with the TE and the allele without the TE for the three replicates analysed. **C**) Schematic representation of the crosses performed to generate two homozygous strains from the Mrp4-TE heterozygous strain T20 (Tokyo): one carrying the insertion (Mrp4-TE+) and the other lacking it (Mrp4-TE-). **D**) Normalized expression of *Mrp4* with *Rp49* in Mrp4-TE+ (red) and Mrp4-TE-(grey) females and males in control (empty boxes) and after paraquat exposure (filled boxes). Boxplot shows the median (horizontal line), first and third quartiles (lower and upper bounds, respectively), and minimum and maximum values (lower and upper whiskers, respectively). Note that the fold changes shown in the figure are based on raw expression values, whereas the fold changes reported in the text are adjusted marginal means from the statistical model, which account for strain and treatment effects. Asterisks indicate level of statistical significance (* p ≤ 0.05, ** p ≤ 0.01, ***p ≤ 0.001).

In females, the mean log₂ ratio of *Mrp4* expression was significantly greater than zero only after paraquat exposure (one-sample t-test: *p* = 0.436 95% CI [-1.241, 1.960], mean = 0.359 for control and *p* = 0.049 95% CI [0.001, 0.688], mean = 0.345 for paraquat; Figure 3B). This indicates that *Mrp4* exhibits allele-specific expression under paraquat-induced stress, with the allele carrying the Mrp4-TE insertion being more expressed. In males, there was a trend toward allele-specific expression under both control and paraquat conditions, with the allele carrying the TE tending to be more highly expressed. However, the mean log₂ ratio did not reach statistical significance, likely due to the small sample size (one-sample t-test: *p* = 0.083 95% CI [-0.176, 1.268], mean = 0.546 for control and *p* = 0.060 95% CI [-0.031, 0.609], mean = 0.289 for paraquat; Figure 3B).

For *inaE*, we could measure expression only in females, as the gene is located on the X chromosome, making it impossible to obtain heterozygous males. We observed allele-specific expression under both control and paraquat conditions, with the mean log₂ ratio of *inaE* expression being significantly greater than zero (one-sample t-test: *p* = 0.029 95% CI [0.011, 0.077], mean = 0.044 for control and *p* = 0.041 95% CI [0.023, 0.447], mean = 0.235 for paraquat; Figure 3B). These results indicate that the allele carrying the inaE-TE insertion was more expressed in both conditions.

Finally, we investigated whether there were polymorphisms linked to the presence of the TE insertion that could also contribute to the observed differences in allele-specific expression. We Sanger-sequenced 1,445 bp of the predicted 3,945 bp coding region of *Mrp4* and identified nine single nucleotide polymorphisms (SNPs), all of which resulted in synonymous substitutions in the predicted protein. Similarly, for *inaE*, we sequenced 1,503 bp of the predicted 2,100 bp coding region and identified 23 SNPs, all producing synonymous substitutions. Thus, although we cannot entirely exclude the contribution of additional polymorphisms to the observed effect, the TE insertions stand out as the most plausible variants driving the changes in gene expression.

Altogether, our ASE experiments suggested that the Mrp4-TE and inaE-TE insertions are likely the causal mutations driving changes in the expression of their nearby genes (Figure 3B). Specifically, the Mrp4-TE insertion was associated with a significant increase in *Mrp4* expression in females following paraquat exposure, with a similar trend observed in males. In contrast, the inaE-TE insertion was associated with a significant increase in *inaE* expression in females under both control and paraquat conditions. These findings are consistent with our observations in wild-type populations, where the presence of Mrp4-TE and inaE-TE insertions appears to enhance paraquat-induced upregulation of their respective genes (Figure 2).

### Mrp4-TE insertion promotes upregulation of *Mrp4* after a paraquat exposure

To validate the effect of the Mrp4-TE on the expression of its nearby gene following paraquat exposure, we generated two homozygous strains from a TE-heterozygous background: one carrying the insertion and the other lacking it. This approach allowed us to directly compare the impact of the TE insertion while minimizing the influence of other polymorphisms (see Materials and Methods). To obtain flies homozygous for the presence or absence of the Mrp4-TE insertion, we started from the heterozygous strain T20 (from Tokyo; Figure 1C, Figure 3C). Individual flies from T20 were crossed, and their offspring were subsequently intercrossed. From these crosses, we identified and selected pairs in which both parents were homozygous either for the presence (Mrp4-TE+) or for the absence (Mrp4-TE-) of the insertion (Figure 3C). Thus, the two strains should be nearly identical at the genomic level, differing only by the presence or absence of the Mrp4-TE insertion or by closely linked polymorphisms, allowing us to attribute any expression differences primarily to the TE itself. Unfortunately, we could not follow the same approach to generate homozygous flies for the presence or absence of the inaE-TE insertion since no heterozygous flies were found among our strains (Figure 1C).

Consistent with our observations in wild-type strains, paraquat exposure affected *Mrp4* expression differently depending on the strain (Figure 3D). In females, Mrp4-TE+ flies showed a significant paraquat-induced increase in *Mrp4* expression (3.3-fold, *p* = 0.011), whereas Mrp4-TE-flies showed no significant change (0.9-fold, *p* = 0.366; Figure 3D). A similar pattern was observed in males: Mrp4-TE+ flies exhibited a significant 3.9-fold upregulation (*p* = 0.004), while Mrp4-TE-flies showed no significant difference (0.3-fold, *p* = 0.791; Figure 3D).

Taken together, these results demonstrate that the Mrp4-TE insertion promotes the upregulation of *Mrp4* expression under oxidative stress conditions.

### Mrp4-TE harbours Cnc binding sites

One possible mechanism by which TEs can enhance the expression of nearby genes is through the presence of TFBSs within their sequences, which may become activated under stress conditions (Chuong et al., 2017; Kunarso et al., 2010; Lynch et al., 2011; Villanueva-Cañas et al., 2019). To test this, we scanned the sequences of *Mrp4* and *inaE*, including their associated TEs and proximal regulatory regions (1 kb upstream of the transcription start site), for the presence of the Cnc transcription factor binding motif, which is implicated in oxidative stress responses. In *Drosophila*, the Cnc-dKeap1 pathway, homologous to the mammalian Nrf2-Keap1 system, has been associated with the regulation of antioxidant and detoxification gene expression (Bayliak et al., 2020; Deng & Kerppola, 2014; Misra et al., 2011; Pitoniak & Bohmann, 2015; Sykiotis & Bohmann, 2008). Indeed, Cnc-dependent activation of target genes has been reported following oral paraquat exposure in *D. melanogaster* (Sykiotis & Bohmann, 2008).

We identified a single Cnc binding site within the Mrp4-TE sequence, whereas no additional sites were detected within the *Mrp4* gene body or its proximal regulatory region (Table S2, Figure S2). However, no Cnc binding sites were found within the inaE-TE sequence, although seven sites were detected within the second intron of *inaE* (Table S2, Figure S2).

Therefore, the association between the presence of Mrp4-TE and the increased expression of *Mrp4* following paraquat exposure may be explained by the presence of a Cnc binding site within the TE sequence, although functional validation will be required to confirm this hypothesis. In contrast, inaE-TE does not seem to contribute additional Cnc binding sites.

### Mrp4-TE is associated with increased sensitivity to oxidative stress

To assess whether the paraquat-induced increase in *Mrp4* expression in flies carrying the Mrp4-TE insertion is associated with an oxidative stress-related phenotype, we performed survival assays after oral paraquat exposure.

In wild-type strains, we found that S29 flies carrying the Mrp4-TE insertion were more sensitive to paraquat than MT47 and W120 flies, which lack the insertion, in both males and females (log-rank test: males, all *p* < 0.001; females, S29-MT47 *p* = 0.035 and S29-W120 *p* < 0.001; Table S4, Figure 4A). Moreover, in males, survival after paraquat exposure did not differ between MT47 and W120 flies (log-rank *p* = 0.620), whereas in females a significant difference was observed between these two strains (log-rank *p* < 0.001; Table S4, Figure 4A). However, in wild-type strains, differences in genetic background may modulate the phenotypic response to paraquat and potentially mask the effect of *Mrp4*. To minimize genetic background variation, we repeated the experiment using Mrp4-TE- and Mrp4-TE+ flies, which differ only by the presence or absence of the TE insertion, respectively. In this controlled genetic context, Mrp4-TE+ males were significantly more sensitive to paraquat than Mrp4- TE- males (log rank *p* = 0.042), and a similar trend was observed in females, although the difference was not statistically significant (log rank *p* = 0.054; Table S5, Figure 4B). To determine whether the same response could be observed with a different oxidative agent, we repeated the survival assays following oral exposure to H₂O₂. Consistent with our previous observations, Mrp4-TE+ flies were more sensitive to H₂O₂ exposure than Mrp4-TE-flies in both males (log-rank *p* = 0.001) and females (log-rank *p* < 0.001; Table S6, Figure S3).

**Figure 4.**
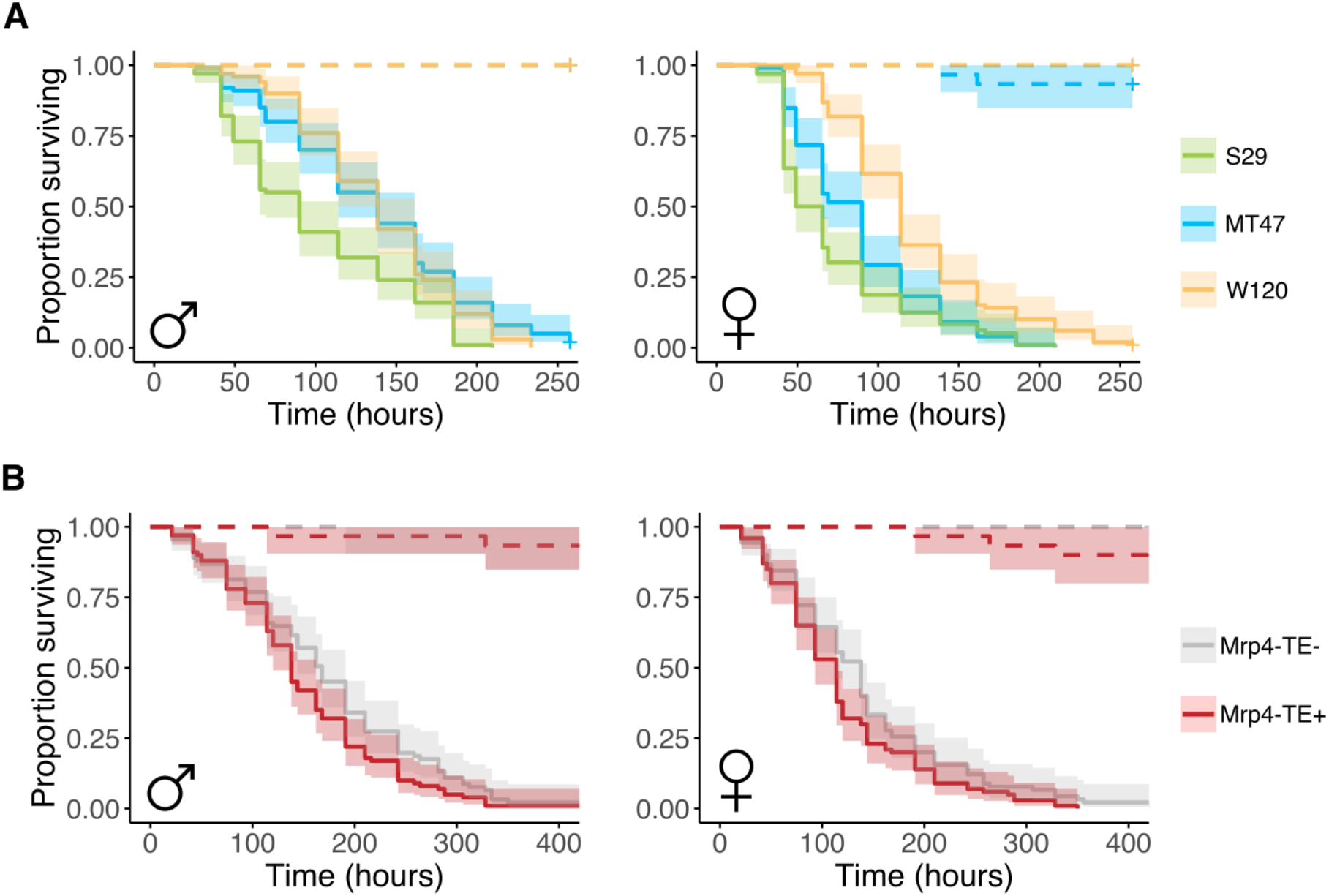
Mrp4-TE insertion in *D. suzukii* may be associated with increased sensitivity to oxidative stress. **A**) Kaplan-Meier survival curves of S29, MT47 and W120 males (left) and females (right) following oral exposure to paraquat. **B**) Kaplan-Meier survival curves of Mrp4-TE+ and Mrp4-TE-males (left) and females (right). The shaded region indicates the upper and lower 95% confidence interval calculated from the Kaplan-Meier curves. Dotted lines represent survival under control conditions, and solid lines represent survival under paraquat exposure.

Overall, our results suggest that the Mrp4-TE insertion in *D. suzukii* may be associated with increased sensitivity to oxidative stress.

## DISCUSSION

In this study, we identified two polymorphic TE insertions, Mrp4-TE and inaE-TE, associated with changes in the expression of their nearby genes in response to oxidative stress in the invasive species *D. suzukii*. We observed that paraquat-induced upregulation of *Mrp4* and *inaE* genes was stronger in wild-type strains carrying these intronic TE insertions. We further validated the regulatory role of both Mrp4-TE and inaE-TE insertions through ASE analyses, which showed that the allele containing the TE exhibited higher expression levels after paraquat exposure. In addition, we found that one of these insertions, Mrp4-TE, contains a binding site for the transcription factor Cnc, a key regulator widely implicated in oxidative stress responses (Bayliak et al., 2020; Deng & Kerppola, 2014; Misra et al., 2011; Pitoniak & Bohmann, 2015; Sykiotis & Bohmann, 2008). Consistent with this observation, analyses of wild-type strains differing only by the presence or absence of the Mrp4-TE insertion revealed a significant paraquat-induced increase in *Mrp4* expression exclusively in flies carrying the insertion.

To our knowledge, these observations provide the first functionally validated evidence in *D. suzukii* of TE insertions associated with gene expression modulation in response to stress. In *D. melanogaster*, several examples of TEs regulating the expression of nearby genes under stress conditions have already been described through gene expression analysis, *in vivo* reporter assays, and allele-specific expression analysis (Guio et al., 2014; Mateo et al., 2014; Merenciano & González, 2023; Ullastres et al., 2021; Villanueva-Cañas et al., 2019). One well-characterized example related with oxidative stress is the TE insertion *Bari-Jheh*, located in the intergenic region of the *Juvenile Hormone Epoxy Hydrolase* (*Jheh*) genes, which has been associated with the upregulation of *Jheh1* and *Jheh2* under oxidative stress induced by paraquat (Guio et al., 2014). In addition to adding antioxidant response elements upstream of *Jheh1* and *Jheh2*, this TE was shown to be enriched for H3K4me3 and H3K27me3 histone marks under oxidative stress conditions, demonstrating that *Bari-Jheh* can modulate the expression of nearby genes both by adding *cis*-regulatory sequences and by altering the local chromatin state (Guio et al., 2014, 2018). Based on our results, we hypothesize that Mrp4-TE promotes the expression of *Mrp4* after paraquat exposure through the addition of a Cnc TFBS. In contrast, we did not identify any Cnc binding sites within the inaE-TE sequence. Therefore, the regulatory effect of this insertion could instead be mediated by the presence of yet unidentified oxidative stress-related transcription factor binding sites or by changes in the chromatin state of the region, although additional experiments will be required to confirm these mechanisms.

In *D. suzukii*, the only previous study investigating the role of TEs in oxidative stress responses in wild-type populations reported a significant depletion of TE insertions near genes differentially expressed after a paraquat exposure, suggesting that purifying selection acts to remove insertions from these loci (Marin et al., 2021). However, our results highlight the importance of studying the effects of individual TE insertions. Although the promotion of oxidative stress-responsive gene expression may not represent a general property of TEs, specific insertions can nonetheless play important roles in shaping transcriptomic responses (Marin et al., 2021). Furthermore, given the high TE content observed in *D. suzukii*, understanding the functional impact of these elements remains particularly relevant.

In this work, we also showed that a *D. melanogaster Mrp4* knockout strain (Mrp4^-/-^) generated using CRISPR/Cas9 genome-editing exhibited almost complete loss of *Mrp4* expression, which was associated with increased sensitivity to paraquat compared with wild-type flies (Huang et al., 2014; Huang & Haddad, 2007). However, in *D. suzukii*, increased *Mrp4* expression after paraquat exposure was not associated with a more resistant phenotype. On the contrary, flies carrying the Mrp4-TE insertion, which displayed higher *Mrp4* expression under oxidative stress conditions likely due to the presence of the TE, appeared to show a slight increase in sensitivity to paraquat and H_2_O_2_. In agreement with our results, a previous study in *D. suzukii* reported differences in survival between two USA strains following acute exposure to broad-spectrum insecticides (Mishra et al., 2018). Interestingly, the strain that was less susceptible to the different insecticide classes showed reduced *Mrp4* expression under control conditions compared with the more susceptible strain (Mishra et al., 2018). Together, these results suggest that increased *Mrp4* expression is not necessarily associated with enhanced resistance to insecticides or paraquat in *D. suzukii*, despite the well-established role of ABC transporters, like the *Mrp4* gene, in detoxification and cellular protection. Nevertheless, additional analyses examining gene expression after insecticide exposure will be necessary to clarify the relationship between *Mrp4* expression and resistance phenotypes.

Another possible explanation for the different relationship between *Mrp4* expression and oxidative stress resistance in *D. melanogaster* and *D. suzukii* is that the effect of *Mrp4* expression may be dosage dependent. While transient induction of *Mrp4* in response to oxidative stress may promote survival during acute stress, constitutive or excessive expression could impose physiological costs or disrupt cellular homeostasis. Consistent with this idea, artificial overexpression of *Mrp4* in *D. melanogaster* has been reported to increase sensitivity to both paraquat- and H_2_O_2_-induced oxidative stress, indicating that higher *Mrp4* expression does not necessarily translate into greater stress resistance (Monnier et al., 2002). Similar dosage-dependent effects have been described for other stress-responsive genes. For example, *Hsp70* is transiently induced by heat stress and enhances thermotolerance, whereas its constitutive overexpression reduces fecundity and larval-to-adult survival in *D. melanogaster* (Krebs & Feder, 1997; Krebs & Holbrook, 2001). We therefore speculate that the increased *Mrp4* expression associated with the Mrp4-TE insertion in *D. suzukii* may exceed the optimal expression level required to produce oxidative stress resistance.

It is also known that conserved detoxification genes can acquire species-specific regulatory roles or display contrasting expression-resistance relationships depending on their ecological and evolutionary context (Berrutti et al., 2026; Harrop et al., 2014; Nguyen et al., 2016). For example, *D. melanogaster* exhibits greater resistance to oxidative stress induced by paraquat or H_2_O_2_ exposure than *D. suzukii* (Nguyen et al., 2016). Comparative analysis of 61 *glutathione-S transferase* (GST) genes revealed that most GST subfamilies present in *D. melanogaster* are also found in *D. suzukii*, although some have not been identified in the latter species yet (Nguyen et al., 2016). Moreover, GST activity assays demonstrated significantly lower overall GST activity in *D. suzukii* (Nguyen et al., 2016). These findings suggest that *D. suzukii* might have evolutionarily adapted to reduce GST activity because its primary food sources are healthy ripening fruits, rather than decaying rotten fruits that release high levels of toxic components (Nguyen et al., 2016). Thus, we hypothesize that, due to the distinct ecological and evolutionary context of *D. suzukii*, *Mrp4* may have acquired a function different from that observed in *D. melanogaster*.

One mechanism underlying species-specific regulatory differences could involve the impact of TEs. In *D. suzukii*, a higher proportion of *cytochrome P450* (CYP) genes involved in detoxification pathways contain TE sequences within or near the genes compared with CYP genes in *D. melanogaster* (Berrutti et al., 2026). This pattern is likely associated with the higher abundance of Helitron insertions in the *D. suzukii* genome (Berrutti et al., 2026; Mérel et al., 2021). Furthermore, CYP-related TE insertions in *D. suzukii* are frequently located in promoter regions, whereas in *D. melanogaster* they are more commonly found within introns, suggesting a putative stronger regulatory role of TEs on detoxification genes in *D. suzukii* (Berrutti et al., 2026). It has also been reported that CYP genes are, on average, 27% longer in *D. suzukii* than in *D. melanogaster*, likely due to the presence of TE insertions (Berrutti et al., 2026). In addition, TEs can induce genomic rearrangements that can result in the duplication of genes, as it has been observed in *D. melanogaster* for genes associated with insecticide resistance (Remnant et al., 2013). Altogether, these findings are consistent with our results and support the idea that TEs may contribute to both gene architecture and gene regulatory evolution in response to environmental pressures.

Finally, from an evolutionary perspective, the low population frequency of Mrp4-TE insertion could be explained by stochastic processes such as genetic drift. Nevertheless, the potentially maladaptive oxidative stress-sensitive phenotype associated with the Mrp4-TE might be another reason explaining its low population frequency. This insertion was found at very low or null frequencies across the three populations analysed (Japan, USA, and France), and was only detected as present or polymorphic in two Japanese strains. However, we cannot exclude the possibility that this TE confers a trade-off effect that has contributed to its maintenance in Japan.

In summary, our results provide the first evidence in *D. suzukii* linking polymorphic TE insertions to changes in gene expression in response to oxidative stress. Furthermore, we found that the TE-associated increase in *Mrp4* expression following paraquat exposure was associated with increased sensitivity to this oxidative agent and to H_2_O_2_. Overall, our work highlights the importance of studying the effects of individual TE insertions in *D. suzukii*, a species whose genome is composed of nearly 50% TE sequences, to better understand the molecular basis of its invasive success. More broadly, our findings also emphasise the importance of investigating gene function within its specific ecological and evolutionary species context.

## DATA AVAILABILITY

All relevant sample information is provided within the manuscript and its Supplementary Materials. All the research complies with applicable laws on sampling from natural populations and animal experimentation (including the ARRIVE guidelines).

## ETHICAL STATEMENT

Ethical approval was not required for this study.

## AUTHOR CONTRIBUTIONS

M.M. and C. V. conceived and designed the study and the experiments. M.M., Z.B., C.R., L.G. and A.C.P.D.D. performed the experiments and analysed the data. M.M. drafted the manuscript. C.V. supervised the project and revised the manuscript. All authors read and approved the final manuscript.

## ACKNOWLEDGEMENTS

We thank the Drosophila Transgenesis Service at the Centro de Biología Molecular Severo Ochoa (Madrid, Spain) for performing the *D. melanogaster* embryo microinjections. We are also grateful to Bénédicte Durand for generously providing *D. melanogaster* Cas9-expressing strains, and to Marta Coronado-Zamora for giving comments on the manuscript. Bioinformatic analyses were performed using the computing facilities of the CC LBBE/PRABI.

