## Supplementary Figures for "Transposable elements modulate gene expression in response to oxidative stress in *Drosophila suzukii*"

**Running title**: TE-associated gene regulation in *D. suzukii*

### SUPPLEMENTARY FIGURES

#### Figure S1

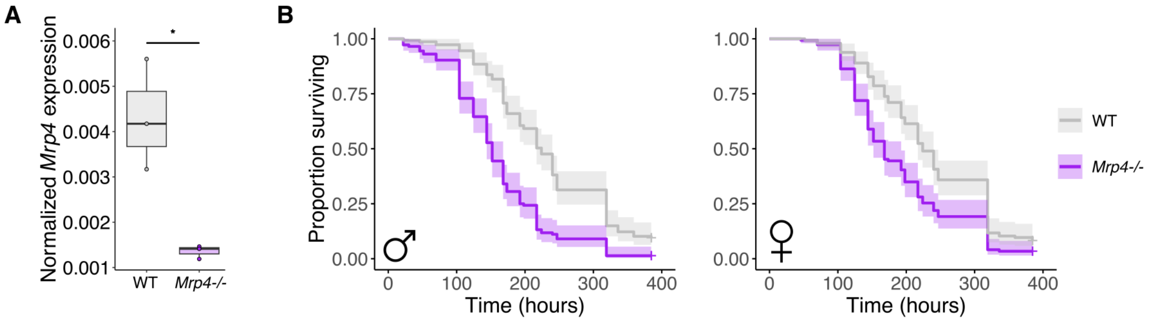


**Figure S1. *Mrp4* expression is involved in promoting resistance to oxidative stress in *D. melanogaster*. A**) Normalized expression of *Mrp4* with *Rpl49* in wild-type (WT) and *Mrp4*$^{-/-}$ flies. Boxplot shows the median (horizontal line), first and third quartiles (lower and upper bounds, respectively), and minimum and maximum values (lower and upper whiskers, respectively). Asterisks indicate level of statistical significance (* $p \leq 0.05$, ** $p \leq 0.01$, ***$p \leq 0.001$). **B**) Kaplan-Meier survival curves of WT and *Mrp4*$^{-/-}$ males and females following oral exposure to paraquat. The shaded region indicates the upper and lower 95% confidence interval calculated from the Kaplan-Meier curves.

### Figure S2

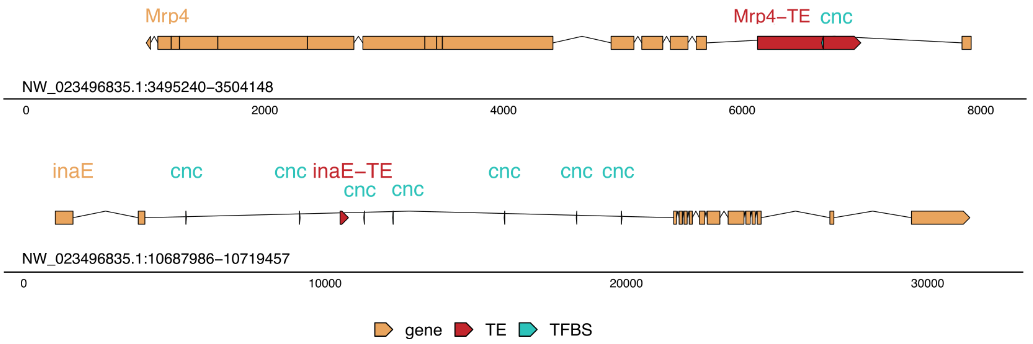


**Figure S2. Schematic representation of the scanned region in the S29 assembled genome with putative Cnc binding sites**. Orange boxes represent gene coding regions and untranslated regions (UTRs) while lines represent introns. TE insertions are represented as red boxes and Cnc binding sites as blue boxes.

**Figure S3**

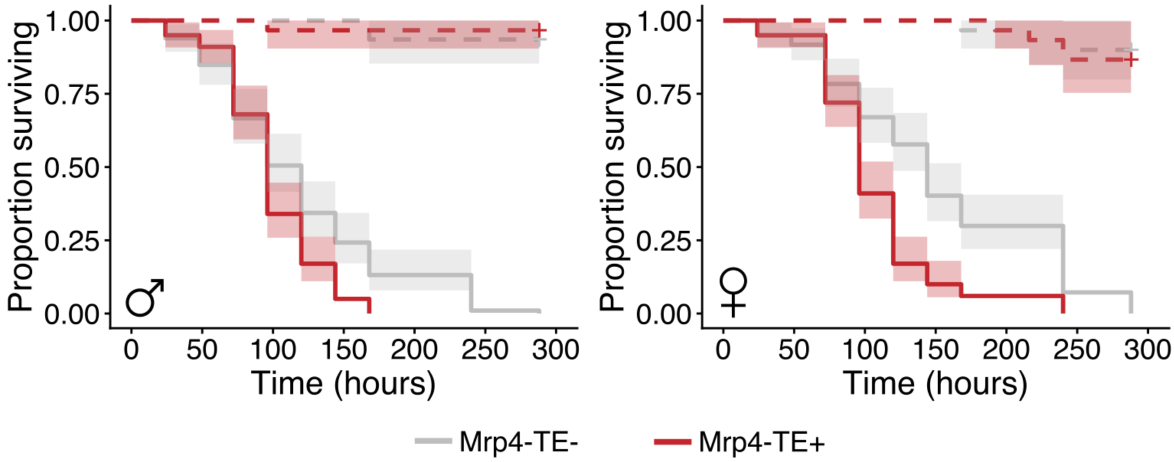


**Figure S3. Survival assay after $H_2O_2$ exposure in Mrp4-TE+ and Mrp4-TE- strains.** Kaplan-Meier survival curves of Mrp4-TE+ and Mrp4-TE- males (left) and females (right). The shaded region indicates the upper and lower 95% confidence interval calculated from the Kaplan-Meier curves. Dotted lines represent survival under control conditions, and solid lines represent survival under $H_2O_2$ exposure.
